# Knockout of *OsSWEET15* impaired rice embryo formation and seed-setting

**DOI:** 10.1101/2022.01.27.478039

**Authors:** Zhenjia Tang, Jing Yang, Shuhui Bao, Zhi Hu, Huihuang Xia, Lai Ma, Qingsong Zheng, Fang Yang, Dechun Zhang, Tai Wang, Shubin Sun, Yibing Hu

## Abstract

we show that the knockout of a sugar transporter gene *OsSWEET15* led to a significant drop in rice fertility because near half of the knockout mutant spikelets bore blighted or empty grains. The rest spikelets bore fertile grains with slightly reduced weight. Notably, the ovaries in the blighted grains of the *ossweet15* mutants expanded after flowering but terminated development before the endosperm cellularization stage and aborted subsequently. GUS and GFP representing OsSWEET15 expression showed that the protein was strongly expressed in the embryo surrounding region (ESR) which was supposed to supply nutrients for the embryo development. These results joined with the protein’s sucrose transport capacity and plasma membrane localization suggest that OsSWEET15 plays a prominent role during the caryopsis formation stage probably by releasing sucrose from the ESR to support the embryo development. By contrast, the empty grains were probably caused by the reduced pollen viability of the *ossweet15* mutants. Investigation of the makeup of *ossweet11* mutant grains revealed similar phenotypes that were observed in the *ossweet15* mutants. These results indicate that both OsSWETT15 and OsSWEET11 play important and similar roles during rice pollen development, caryopsis formation, and seed-setting in addition to their function in seed-filling that was demonstrated previously.

---

Pollen development and pollen tube growth are identified to be important to seed-setting rate and product (Higashiyama and Takeuchi, 2015; Arshad et al., 2017; Xu et al.,2017). By contrast, other developmental factors that affect the seed-setting rate and their mechanisms have not been fully addressed. In the seeds of *Arabidopsis* and cereal crops, the embryo is surrounded by the micropylar endosperm (MCE) and the embryo surrounding region (ESR), respectively (Olsen, 2001; 2004; Berger, 2003). Accumulating data show that the embryo is symplastically isolated from the endosperm shortly after the embryo differentiation due to a lack of plasmodesma between the embryo and ESR/MCE in angiosperms (Suzuki et al., 1993; Demanson, 1997; Opsahl-Ferstad et al., 1997; Kawashima and Goldberg, 2009; Lafon-Placette and Köhler, 2014). The embryo thus takes in nutrients from the MCE/ESR-embryo interface by itself (Morley-Smith et al., 2008; Lafon-Placette and Köhler,2014). These nutrients must be first released from the MCE/ESR into the interface (apoplast) before being actively or passively transported into the embryo.

Among the nutrients necessary for embryo development, sugar is by far the most abundant component. Previous research shows that sucrose-proton symporters (SUC/T) play important roles in the uptake of sucrose from the apoplast into cytoplasm across the membrane of the cells at the acceptor (e.g., embryo) side (Matsukura et al., 2000; Zhang et al., 2007; Bai et al., 2015; Wang et al., 2021). How sucrose is released from the donor cells (e.g., endosperm) on the other side of the interface remains a question until *SWEET* gene is characterized (Chen et al., 2010; Eom et al., 2015). SWEETs are sugar transporters that translocate mono-/disaccharide sugars across the membrane along their concentration gradients. Some of them are also capable of transporting GA in *Arabidopsis* and rice (Kanno et al., 2016; Morii et al., 2020). Physiological functions of SWEETs in model plants include phloem loading and unloading, nectar secretion, transport sugar between organelles, and providing sugar for symbiont or pathogen propagation (Chen et al., 2012; 2015a;b; Guo et al., 2014; Lin et al., 2014; Sosso et al., 2015; Li et al., 2017; Oliva et al., 2019; Ko et al., 2021). At the molecular level, they play a critical role in the intercellular sugar transport when cells lack plasmodesma connection (Chen et al., 2015b).

In rice, besides the interface between the embryo and its surrounding ESR, another interface in caryopsis that lacks plasmodesma connection is the apoplast between the nucellar epidermis and aleurone. By CRISPR-Cas9 or TALEN mediated knockout strategies, Ma et al. (2017) and Yang et al. (2018) demonstrated that the nucellar epidermis-abundantly expressed OsSWEET11 plays an essential role in the early seed-filling stage by acting as a sucrose transporter to release sucrose from the nucellar epidermis; and knockout of the gene impaired grain-filling, reduced seed-setting rate, and prolonged seed maturation time of rice (Ma et al.,2017; Yang et al.,2018; Fei et al., 2021). Moreover, Yang et al. (2018) reported that OsSWEET15 took part in the seed-filling process, either; since double mutation of *OsSWEET11* and *OsSWEET15* completely sterilized the mutant rice, but knockout of *OsSWEET15* alone did not confer any abnormal phenotype. Despite that these investigations answered how sugar is released from the nucellar epidermis to the maternal-filial interface for rice endosperm development, how sugar is released from the ESR for the embryo development during rice caryopsis formation is still not clear.

Here, we report that OsSWEET15, a sucrose transporter (Yang et al.,2018), was strongly expressed in rice caryopsis particularly in the ESR after flowering in addition to its expression in the integument, nucellar epidermis, pollen, and other tissues. Knockout of *OsSWEET15* via CRISPR-Cas9 mediated gene editing led to a significant drop in rice seed-setting rate because only about half of the mutant spikelets bore fertile seeds. Inspection of the infertile spikelets revealed that about 40 % of their caryopses terminated development before the endosperm cellularization stage. These caryopses gradually withered and their spikelets developed into blighted grains. About 60 % of the infertile spikelets did not show ovary expansion after flowering and formed empty grains. Moreover, the 1000-grain weight of the fertile grains of the *ossweet15* mutants reduced slightly compared with that of the wild type (WT) control. As GUS and GFP representing gene expression consistently indicate that OsSWEET15 was expressed in the embryo-surrounding ESR which is supposed to be the nutrient supplier for the embryo development, taking the sucrose transport capacity of OsSWEET15 and its plasma membrane localization tested in rice protoplast into account, we proposed that the caryopsis abortion of the *ossweet15* mutants is likely due to the deficient supply of sugar during the embryo development. Moreover, the empty grains were likely caused by the reduced pollen viability of the *ossweet15* mutants. Observation of *ossweet11* mutants revealed similar phenotypes.

These results imply that both OsWEET15 and OsSWEET11 play important roles in rice pollen function, caryopsis formation and seed-setting, in addition to their function in seed-filling which was reported previously (Ma et al.,2017; Yang et al.,2018; Fei et al., 2021).

## RESULTS

### Significant Reduction of Seed-Setting Rate of the *OsSWEET15* Knockout Mutants

Among the CRISPR-Cas9 edited *OsSWEET15* mutant lines based on rice cultivar Nipponbare (*Oryza sativa* L. ssp. *japonica*), many suffered from marked decreases in their seed-setting rates compared with WT control (Fig 1). Therefore, three independent transgenic lines, *m15-5, m15-9*, and *m15-27* were selected for further investigation (*m15-5* and *m15-9* derived from Spacer 2 and *m15-27* from Spacer 1, both spacers targeted the third exon of the gene, spacer sequences were listed in Supplemental Table S1). Both the *m15-5* and *m15-27* contain more than one nucleotide deletion, while the *m15-9* contains a single nucleotide insertion in the target region of *OsSWEET15* (Supplemental Fig. S1). The mutated genes either encode truncated proteins or amino acid-missing protein (Supplemental Fig. S2). The deletions and insertion located in the target DNA regions (Supplemental Fig. S1), and fragment amplification of the mutant lines for verification of the gene-editing was performed using a high-fidelity polymerase (PrimeSTAR, Takana).

**Figure 1.**
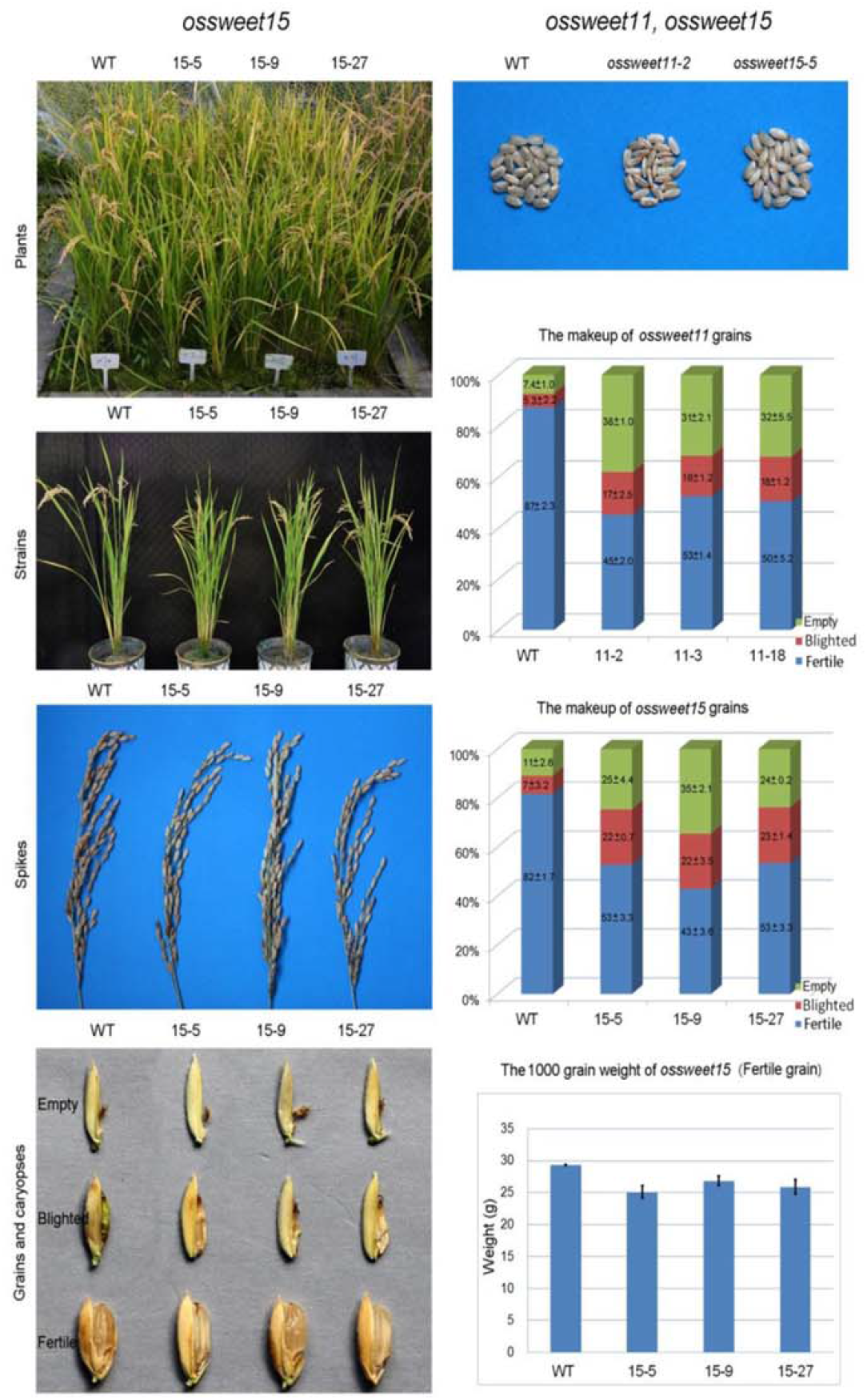
Comparisons of the phenotypes of *ossweet15, ossweet11*, and WT control. The left column shows plants, strains, spikes, grains, and caryopses of *ossweet15* mutants and WT of rice cultivar Nipponbare. The right column shows the images of caryopses, statistical analyses of caryopsis makeups of *ossweet15, ossweet11*, and the 1000-grain weight of *ossweet15*.

As Yang et al. (2018) reported they did not observe any abnormal phenotype in their knockout mutants of *OsSWEET15* based on rice cultivar Kitaake (*Oryza sativa* L. ssp. *japonica*), we first checked the specificity of the gene-editing effect in our investigation. Nucleotide sequences partly identical to the spacers particularly at the PAM end from the genome of Japonica rice Nipponbare were retrieved and examined because they might interfere with the specificity of the CRISPR-Cas9 mediated gene-editing (Jinek et al., 2012). The spacer 1 shares 14 bp nucleotides at most with 3 additional genomic DNA fragments and it shares 13 bp nucleotides with 4 additional genomic DNA fragments other than its target sequence (Supplemental Fig. S3). All these partly identical sequences locate at the interval regions far from the putative promoter or coding region of the predicted genes they adjoin and only two of them possess the critical PAM (NGG) sequences (Supplemental Fig. S3). The spacer 2 shares 13 bp nucleotides at most with additional 133 genomic DNA fragments other than its target sequence. As mentioned above, many lines from the CRISPR-Cas9 plasmids-mediated transgenic rice showed consistent decreases in their seed-setting. It is unlikely that both of the spacers confer similar phenotypes even if spacer 2 aims at non-target genes by error. We thus reasoned that the phenotype we observed in the *OsSWEET15* transgenic rice was not an off-targeting effect.

To exclude the possibility that the phenotypes we observed in the *OsSWEET15* mutants were the consequences of T-DNA’s random insertions into functional gene locus in the rice genome, we introduced the CRISPR-Cas9 gene-editing plasmids into Nipponbare callus for the second time (performed in 2021), and obtained similar results. From the year 2017, we have grown and checked the phenotypes of the *ossweet15* mutants for 6 rounds in 5 years. Particularly, we checked the *ossweet15* mutant’s genotypes once again in 2021 and confirmed their identities (data not shown). The *ossweet15* mutants showed consistent phenotypes except that their seed-setting rates fluctuated to some extent due to yearly weather changes. Taking all the facts into account, we conclude that the reduction of seed-setting rate in the transgenic rice that we observed was caused by the knockout of *OsSWEET15*.

Detailed inspection revealed that the *ossweet15* mutant rice lines did not show obvious differences at the vegetative growth stage. At the reproductive growth stage, however, the mutant spikes were lighter than that of the WT rice because near half of their spikelets bore blighted grains or were simply empty (Fig. 1). The seed-setting rate of the WT control was about 82 % based on statistics investigated in 2021 with at least 800 grains in each sample. By contrast, for the three mutant lines of *m15-5, m15-9*, and *m15-27*, their seed-setting rates were approximate 53 %, 43 %, and 53 %, respectively (Fig.1). Observation of the spikelets of the three mutant lines showed that averagely about 22 % of them formed blighted grains (Fig. 1); and about 28 % of the spikelets did not show any sign of ovary development after flowering, they thus developed into empty grains. The rest spikelets (about 50 %) seemed normally developed and bore fertile grains (Fig.1). However, the 1000-grain weights of the fertile grains reduced slightly compared with that of the WT control (Fig.1). Notably, the WT rice also bore blighted grains and empty grains, but their makeups were about 7 % and 11 %, respectively.

Previously, Ma et al., (2017) and Yang et al. (2018) demonstrated that knockout of *OsSWEET11* impaired seed-filling of the transgenic rice. Moreover, Ma et al. observed that the seed-setting rate of the *ossweet11* mutants also reduced (Ma et al., 2017). This phenotype was consistent with the investigations of Chu et al. (2006) and Yang et al. (2006) based on RNAi-mediated knockdown rice lines of the gene. As the expressions of *OsSWEET11* and *OsSWEET15* largely keep pace with each other according to RiceXPro (https://ricexpro.dna.affrc.go.jp/) and Yang et al. (2018) who showed that OsSWEET11 and OsSWEET15 have a very similar expressional pattern in caryopsis, we thus checked the makeup of the *ossweet11* mutant grains and found that similar to the observation that we obtained in the *ossweet15* mutants, the percentages of fertile grain, blighted grain, and empty grains of the *ossweet11* mutants (*11-2; 11-3; 11-18*. Ma et al., 2017) were approximately 49 %, 17 %, and 34 %. By contrast, the percentages in the WT rice were 87 %, 5 %, and 8 %, respectively (Fig.1; Supplemental Fig. S4).

### Caryopsis Abortion in the Blighted Grains of the *ossweet15* Mutants

To explore the underlying reason for the reduction of seed-setting rate, we first compared floret morphology and structure of the *ossweet15* mutants and the WT control but did not find any obvious difference (Fig. 2). KI-I_2_ staining and germination test showed that both the pollen viabilities and pollen germination rates of the *ossweet15* mutants reduced compared with that of the WT control (Fig. 2). These results partly explained that the seed-setting rates of *ossweet15* mutant lines were significantly lower than that of the WT control.

**Figure 2.**
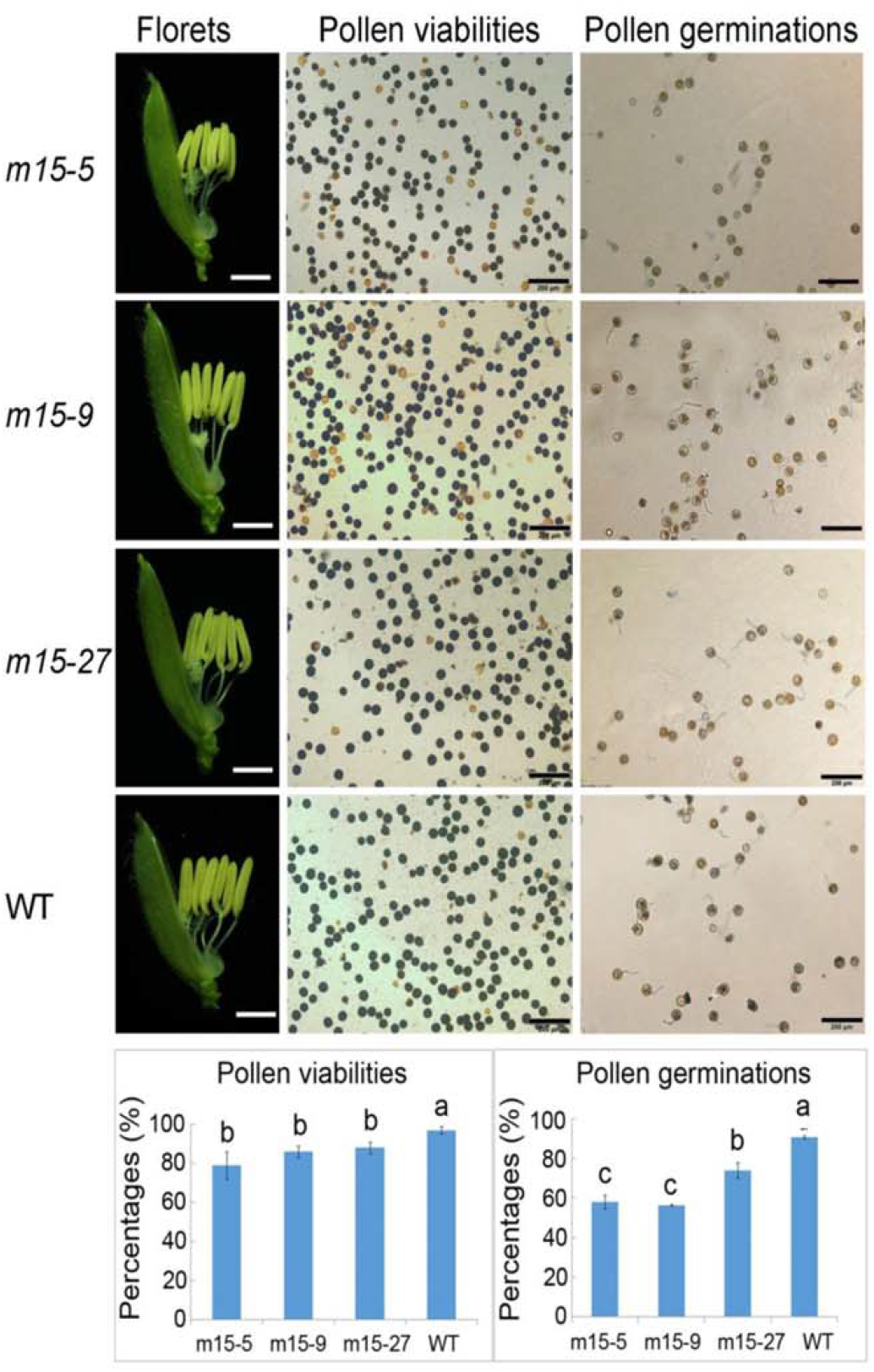
Comparisons of the *ossweet15* mutant lines (*m15-5; m15-9; m15-27*) and WT rice. The first lane shows florets, bars=1 mm; the second lane shows pollens stained with KI-I_2_ solulion; the third lane shows pollen germination on a medium observed under a microscope (Olympus BX 51), bars=200 μm in the second and third lanes. The columns at the bottom of the figure are pollen viabilities and germination rates of *ossweet15* mutant lines and WT rice as control.

Examining the caryopsis development of the *ossweet15* mutants and WT control further revealed their differences (Fig. 3). At 3 days after flowering (DAF), about half of the *ossweet15* mutant’s spikelets developed normally and approximately one-quarter of them bore slightly slimmer caryopses compared with that of the WT control. The rest failed to develop because their ovaries did not show any sign of expansion. At 5 DAF, differences in size between the slimmer and normal caryopses became clear since the development of the slimmer caryopses seemed terminated. The spikelets with the unexpanded ovaries remained undeveloped. At 9 DAF, the normally developed caryopses of the mutants and most caryopses of the WT control reached their full sizes. Longitudinal sections of their caryopses showed that their embryos had fully developed and their endosperms had solidified due to starch-formation (Fig. 3). By contrast, the development-terminated caryopses of the mutant lines began to wither. Longitudinal sections of them showed that the embryo sacs were still filled with clear sap. Notably, all of the development-terminated caryopses lack the embryo (Fig. 3). At 15 DAF, when WT caryopses and normally developed caryopses of the mutants entered into the dough stage, the development-terminated caryopses of the mutant lines aborted and withered gradually. The spikelets with the aborted caryopses developed into blighted grains. It is worth noting that knockout of *GmSWEET15* and *SlSWEET15*, the homolog genes of *OsSWEET15* in soybean and tomato, confer severe seed abortion according to the investigations of Wang et al. (2019) and Ko et al. (2021).

**Figure 3.**
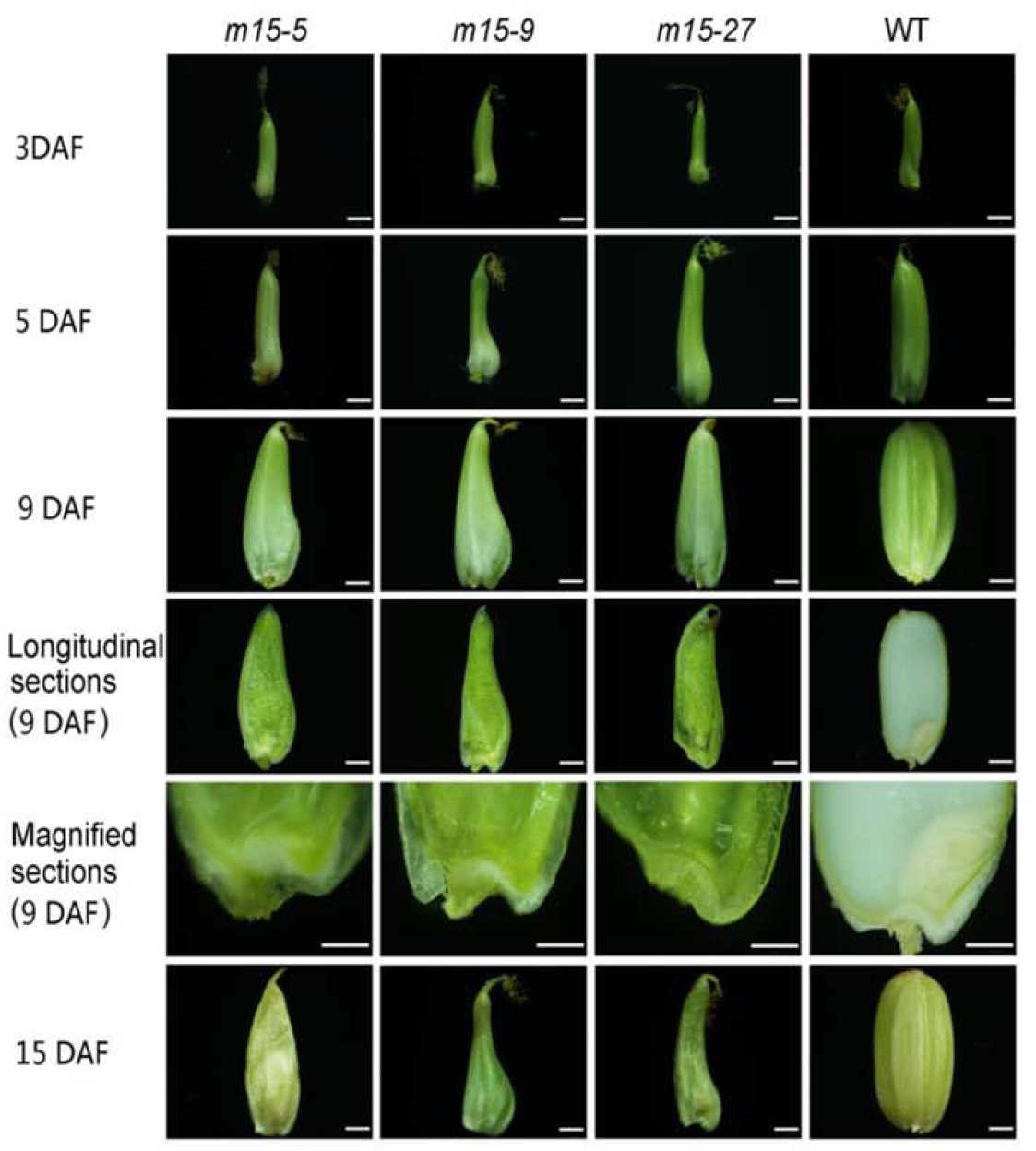
Morphology and structure of the development-terminated caryopses (Withered) of *ossweet15* mutant lines (*m15-5; m15-9; m15-27*) and WT rice (Nipponbare) at different developing stages. DAF: days after pollination. Bars=1 mm.

### Failure of Embryo Formation at the Endosperm Cellularization Stage in the Aborted Caryopsis of the *OsSWEET15* Mutants

In angiosperms, the endosperm development can be divided into the free nuclear stage (coenocyte and alveolar) characterized by nucleus division without cell wall formation, follows by the cellularization stage in which endosperm cell wall established and endosperm cells become independent from each other (Olsen, 2004). These two stages in rice cultivar Nipponbare last about 2 and 2-3 days, respectively (Wu et al., 2016a, b). As the endosperms of the aborted caryopses failed to cellularize, saps were sucked out from the embryo sacs of these caryopses and WT control at 4 DAF and were stained with 4’,6-diamidino-2-phenylindole (DAPI) solution. The result showed that the nuclei were still in a free state in the development-terminated caryopses of the blighted grains (Fig. 4, upper panel). By contrast, the nuclei were wrapped with cytoplasm materials (Fig. 4, upper panel) in most of the WT grains and the normally developed mutant grains. Moreover, the peripheral parts of the embryo sac in the normally developed caryopses have cellularized and solidified. These results indicate that the free nuclear stage was normal but the subsequent cellularization stage was impeded in the aborted caryopsis. Traverse sections of *m15-9* caryopses at 4 DAF showed that no embryo emerged in the development-terminated caryopses (Fig. 4, lower panel, left). By contrast, the embryo had established at a similar site in the normally developed caryopsis at 3 DAF (Fig. 4, lower panel, right).

**Figure 4.**
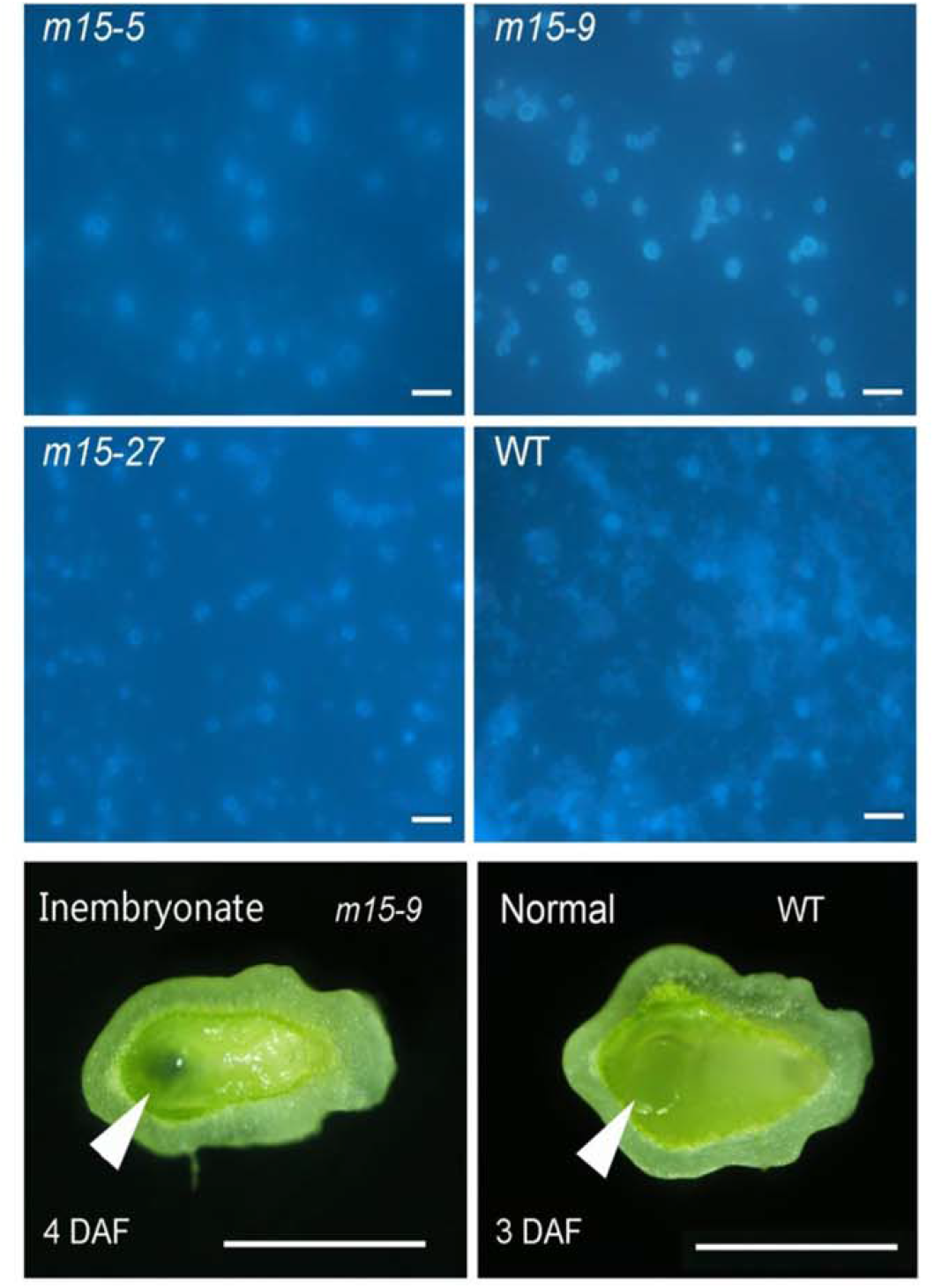
Comparisons of the embryo and endosperm of *ossweet15* mutants and WT rice at the early stage of rice caryopsis development. Upper panel: fluorescence observation of syncytium (multinucleate cell) in the embryo sac of *ossweet15* mutant lines (*m15-5; m15-9; m15-27*) and WT rice at 4 DAE Saps that were sucked out from the embryo sacs of the mutants and WT rice were stained with a DAPI solution before being observed under a microscope (Leica DM5000B). Lower panel: traverse sections of the develop-stagnated caryopsis (4 DAF) and the normally developed caryopsis (3 DAF) near the bottom of seeds in the *m15-9* mutant rice line. Bars=10 μm in the upper panel, bars=200 μm in the lower panel.

### Strong Expression of OsSWEET15 in the Embryo Surrounding Region of Rice Caryopsis

qRT-PCR result (Fig. 5 A) showed that *OsSWEET15* was strongly expressed in anthers and caryopses of rice during the early stages of development after flowering. Moreover, it was weakly expressed in the basal stem or root and leaf. This expression pattern was consistent with the Affymetrix chip-based gene expression profile (http://ricexpro.dna.affrc.go.jp/) which shows that *OsSWEET15* was expressed very abundantly in the ovary and endosperm. Histochemical analysis of pOsSWEET15::GUS expression (Fig. 5B, C, F-H) and observation of pOsSWEET15::GFP fluorescence (Fig. 5D, E) in caryopses at 5 DAF from at least 3 independent transgenic lines consistently showed that *OsSWEET15* promoter activity was evident in rice caryopsis, and especially strong in the ESR (Fig. 5B-H) including the nucellar epidermis and aleurone cells adjoining the embryo (Fig. 5F,G). It was also strongly expressed in the cross cell, integument, and embryonic epidermis (Fig. 5G). Moreover, *OsSWEET15* promoter activity can be found in pollen (Fig. 5H), root, leaf blade, sheath, and stem (Supplemental Fig. S5). These results suggest that OsSWEET15 plays multiple roles in rice development besides its important functions in pollen development and caryopsis formation and filling.

**Figure 5.**
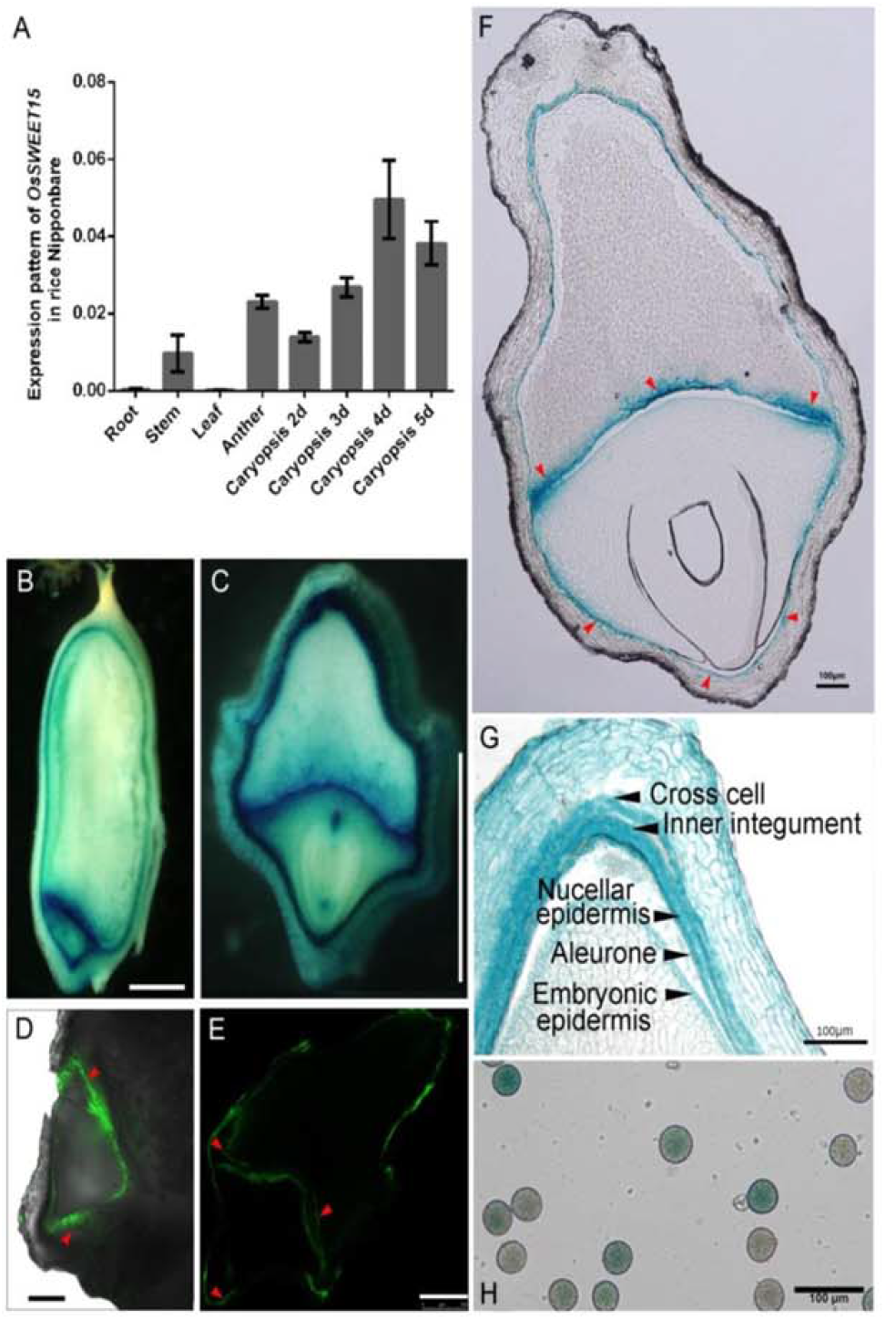
Expressional profiles of *OsSWEET15* in wild type (WT) rice Nipponbare at the transcriptional level and translational level. A: transcriptional levels of *OsSWEET15* in rice tissues examined by real-time qRT-PCR. Caryopsis 2d, 3d, 4d, and 5d show caryopsis samples collected on the 2, 3. 4, 5 DAF. Root, stem, and leaf were collected from 3 week-old rice plants; B, C: longitudinal and traverse sections of rice caryopses expressing pOsSWEETI5::GUS at 5 DAF after GUS staining; D, E: GFP fluorescence observation of longitudinal and traverse sections of rice caryopses expressing pOsSWEET15::GFP at 5 DAF; F: a traverse section cross the embryo area of the rice caryopsis expressing pOsSWEET15::GUS at 5 DAF. The image was observed under a microscope (Leica, DM5000B); G: a traverse-sectioned image of the rice caryopsis expressing pOsSWEET15::GUS at 5 DAF to show the detailed tissue organizations. H: rice pollens expressing pOsSWEET15::GUS observed under a microscope (Olympus BX51) after GUS staining. In B-F, En: endosperm; Em: embryo. Red arrowheads in D, E, F indicate the ESR. Bars in B and C =1 mm; in D and E =250 μm, in F, G, and H =100 μm.

### Subcellular Localization of OsSWEET15-EGFP Fusion Protein

OsSWEET15 and EGFP fusion protein was used to indicate the subcellular localization of OsSWEET15 in rice protoplast. A membrane protein AtPIP2;1 fused with mCherry was employed as the plasma membrane marker (Breia et al., 2020). Different from OsSWEET11-EGFP fusion protein located on the plasma membrane (Fig. 6; Ma et al., 2017), OsSWEET15-EGFP fusion protein is also expressed in the cytoplasm in addition to expressing on the plasma membrane of the protoplast. We repeated the experiment 3 times and obtained consistent results.

**Figure 6.**
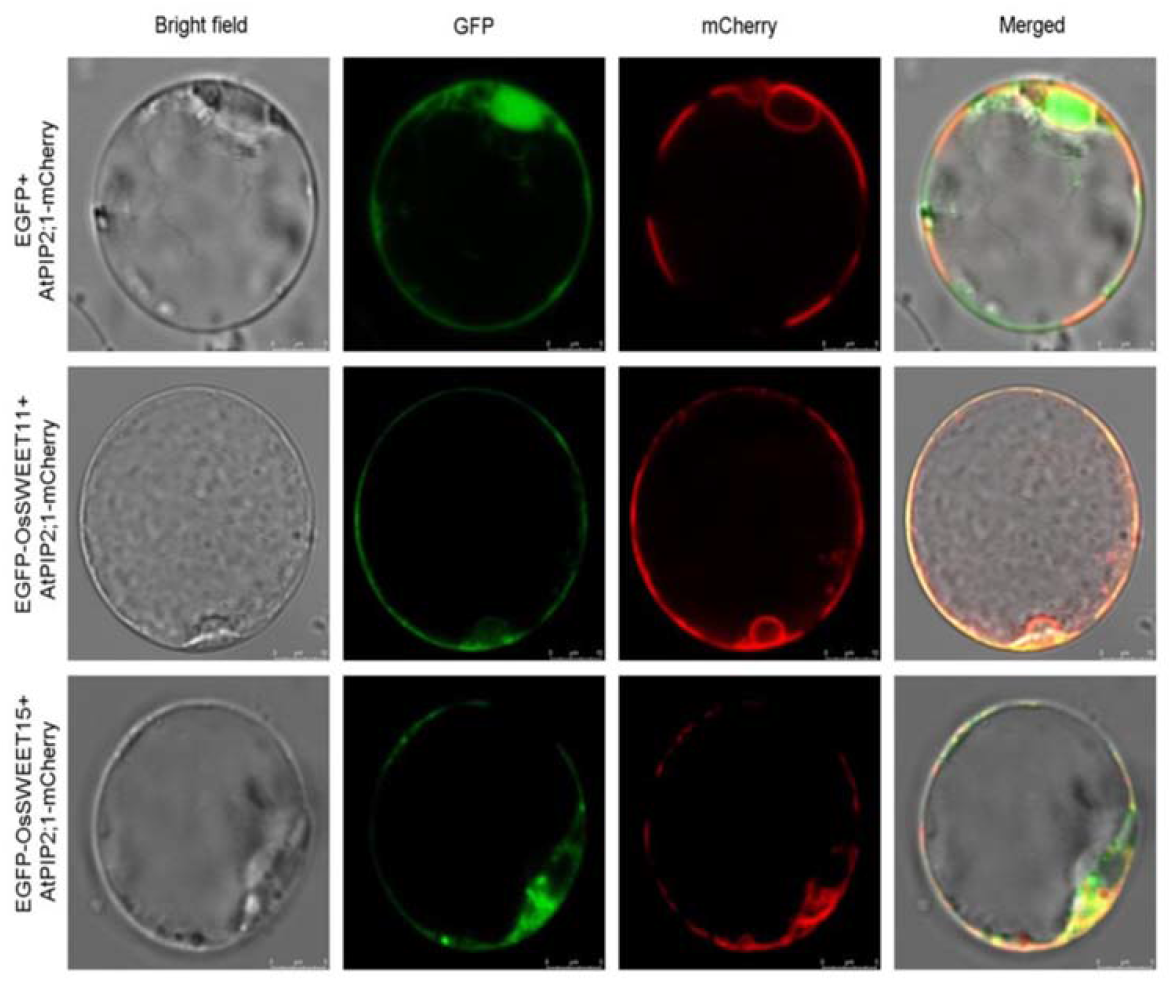
Subcellular localization of OsSWEET15–EGFP fusion protein in rice protoplasts. Plasmids consisting of pSAT6A–OsSWEET15-EGFP and CD3-1007 were co-transfected via a PEG-mediated method. Plasmids consisting of pSAT6A–EGFP and CD3-1007 were co-transfected in parallel as a negative control and plasmids consisting of pSAT6A–OsSWEET11-EGFP and CD3-1007 were co-transfected as a positive control. EGFP fluorescence was observed under a Leica SP5 confocal microscope. Bars in the first and the third rows =10 μm; bars in the second rows=5μm.

## DISCUSSION

Rice caryopsis development can be divided into three consecutive stages including seed (caryopsis) formation, seed filling, and seed maturation. We show that the knockout of *OsSWEET15* led to about 22 % of their caryopses terminated development at the embryo formation stage and eventually forming blighted grains (Fig. 3, 4). In addition, about 28 % of spikelet’s ovaries failed to develop and formed empty grains suggesting that the protein functioned in pollen development and probably also in the fertilization process (Fig. 2). Moreover, the 1000-grain weight of the fertile grains of the *ossweet15* mutants decreased slightly (Fig. 1) and double mutation of *OsSWEET11* and *OsSWEET15* led to completely infertile (Yang et al., 2018; Supplemental Fig. S6) implies that OsSWEET15 participated in the seed-filling process as well.

Endosperm cellularization represents an important developmental transition during seed development (Olsen, 2001; Hehenberger et al., 2012). It initiates at the embryo vicinity and follows by the peripheral endosperm region (Engell, 1989; Olsen, 2001). Finally, cellularization takes place in the center region of the endosperm (Suzuki et al., 2000; Itoh et al., 2005; Lafon-Placette and Köhler, 2014). Ultrastructural observation in maize, soybean, *Arabidopsis* and *Solanum nigrum* revealed that the embryo is symplastically isolated from the surrounding endosperm shortly after the embryo cell differentiation (Schel et al., 1984; Dute et al. 1989; Mansfield and Briarty, 1991; Briggs, 1996). Analogously, Suzuki et al. (1993) reported that no plasmodesma was observed between the embryo and the endosperm cells in rice caryopsis at 42 hours after pollination. We checked the caryopses of rice cultivar Nipponbare at 5 DAF and confirmed Suzuki’s observation although there are many plasmodesmata between the cells within the embryo or endosperm (Fig. 7). These facts imply that nutrients in or out of the embryo after ESR cellularization have to be transported across the plasma membranes via transporters.

**Figure 7.**
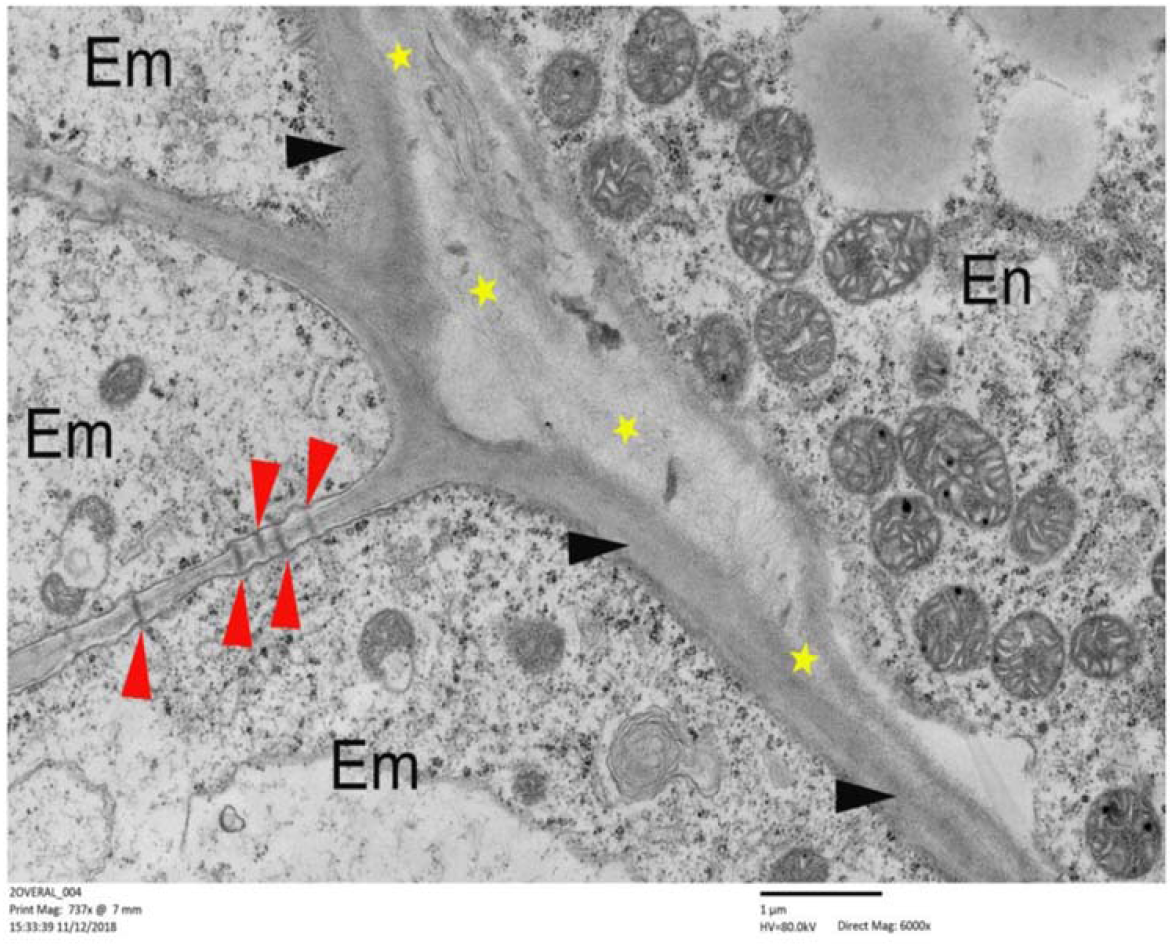
Observation of a longitudinal section of the embryo and endosperm in WT rice caryopsis at 5 DAF under a transmission electron microscope (Hitachi H-7650). Yellow asterisks indicate the interface of the embryo and endosperm. Red arrowheads indicate plasmodesmata. Em: embryo cell; En: endosperm cell. Bar=2 μm. Magnification rate=3000.

GUS and GFP representing protein expression showed that OsSWEET15 was strongly expressed in the ESR region (Fig. 5). This expression pattern is similar to the reports that AtSWEET15 and GmSWEET15 were expressed in the micropylar endosperms of *Arabidopsis* and soybean seeds (Chen et al., 2015b; Wang et al., 2019). It is also consistent with the report that SlSWEET15 was expressed in the tomato seed coat encompassing the embryo (Ko et al., 2021).

MCE/ESR is supposed to be the tissue responsible for nutrient supply for embryo development (Olsen, 2001; 2004; Dumas and Rogowsky, 2008). Taking the sucrose transport capacity (Yang et al., 2018) and plasma membrane localization of OsSWEET15 (Fig. 6), particularly that about 22 % of the *ossweet15* mutant caryopses lacked the embryos and they aborted at the endosperm cellularization stage (Fig. 4) into account, we propose that OsSWEET15 played an important role in the seed formation stage likely by providing sucrose for the embryo development (Fig 5F, G). Therefore, the knockout of *OsSWEET15* led to the termination of embryo development since the ESR is essential for embryo-endosperm communication (Becraft, 2001; Olsen et al., 2004; Zhou et al., 2013).

Previously, both Ma et al (2017) and Yang et al. (2018) reported that the knockout of *OsSWEET11* affected the seed-filling of rice. In addition, Ma et al. (2017) reported that the seed-setting rate of *ossweet11* mutants reduced and Yang et al. (2018) reported that the maturation of *ossweet11* seeds was prolonged. We scrutinized the *ossweet11* mutants based on rice cultivar Nipponbare (in 2021) and confirmed these phenotypes. However, Yang et al. (2018) reported that mutation of *OsSWEET15* alone did not confer any abnormal phenotype whereas our data showed clearly that OsSWEET15 exerted significant impacts on rice seed-setting rate and pollen viability (Fig.1–4). Notably, both Wang et al. (2019) and Ko et al. (2021) demonstrated that the CRISPR-Cas9 mediated knockout of *GmSWEET15* and *SlSWEET15*, the homolog genes of *OsSWEET15* in soybean and tomato, led to serious defects in seed-filling and embryo development. As a result, more than 80 % of soybean seeds and most tomato seeds were aborted in the genes mutants. These phenotypes are very similar but more severe than what we observed in our *ossweet15* mutants. Wang et al. (2019) and Ko et al. (2021) concluded that GmSWEET15 and SlSWEET15 played critical roles in sucrose unloading from the endosperm or seed coat to support the developing embryos. These conclusions support our proposal that OsSWEET15 releases sucrose from the ESR including nucellar epidermis and aleurone to nurture the embryo’s development in rice.

It should be mentioned that both Yang et al. (2018) and our results (Fig. 5F) showed that OsSWEET15 was strongly expressed in the nucellar epidermis of rice caryopsis. However, we also showed that the protein was strongly expressed in the ESR region as we traverse-sectioned rice caryopsis cross the embryo proper. One possible reason for the discrepancy between Yang et al. (2018) and our results (Fig. 1–4) is that different gene-editing methods (Yang et al. (2018) used TALEN while we used CRISPR-CAs9) or/and different rice cultivars (they use Kitaake while we used Nipponbare) were used for the investigations. Actually, the impairment in the seed-filling of the *ossweet11* mutants reported by Yang et al. (2018) is much milder than that we found in our *ossweet11* mutants (2017) judged from the images and descriptions in their article.

Consistent with the results of Chu et al. (2006) and Yang et al. (2006) based on their RNAi knockdown rice lines, our results showed that knockout of *OsSWEET11* reduced rice seed-setting rate (Ma et al., 2017; Fig.1, Supplemental Fig. S4). As *OsSWEET11* and *OsSWEET15* have a similar expression pattern in the developing caryopsis (RiceXPro https://ricexpro.dna.affrc.go.jp/; Fig. 5F, Ma et al., 2017; Yang et al., 2018), it is not surprising that the makeups of fertile grain, blighted grain, and empty grain of *ossweet11* and *ossweet15* mutant were similar (Fig.1; Supplemental Fig. S4). Moreover, knockout of *OsSWEET11* and *OsSWEET15* that strongly expressed in the nucellar epidermis (Fig. 5F, Ma et al.,2017; Yang et al., 2018) reduced rice grain weight similarly but in varying degrees (Ma et al., 2017; Fig.1). Double mutation of *OsSWEET15* and *OsSWEET11* sterilized the transgenic rice (Supplemental Fig. S6; Yang et al., 2018). These results indicate that both of them play important roles in rice seed-filling.

Interestingly, our results showed that OsSWEET15-EGFP was also expressed in the cytoplasm of rice protoplast in addition to the plasma membrane. Moreover, the fluorescence of the OsSWEET15-EGFP fusion protein was weaker than that of the OsSWEET11-EGFP fusion protein under the same cultivation condition. Future investigation may reveal the underlying reason.

## MATERIALS AND METHODS

### Gene Cloning and Plasmid Preparation

The coding sequence of *OsSWEET15* (LOC_Os02g30910) was amplified using rice inflorescence cDNA template and P1/P2 primers (sequences listed in Supp. Table 1) based on a full-length cDNA clone (accession No. AK103266). The amplified fragment was then cloned into a pEASY-Blunt vector (Transgene Biotech) and sequenced. A putative promoter of *OsSWEET15* (genomic sequence of *OsSWEET15* from the translation start code ATG to upstream 2.262 Kb) was amplified with rice Nipponbare genomic DNA template and primers P3/P4 with restriction sites *Pst*I and *Bam*HI added, respectively. It was integrated into a modified pCAMBIA1300 vector (contains a GUS-coding sequence) or pCAMBIA1302 to drive GUS or GFP expression (pOsSWEET15::GUS or pOsSWEET15::GFP). Open reading frame (ORF) sequence of *OsSWEET15* was amplified with P5/P6 (with restriction sites *Hin*dIII and *Bam*HI) and P7/P8 primers (with *Hin*dIII and *Bam*HI). They were integrated into pSAT6A-EGFP-N1 and pSAT6A-EGFP-C1 (Chung et al. 2005), respectively, in-frame with the EGFP-coding sequence to create fusion proteins after digestion. Subsequently, the plasmids were digested with restriction endonuclease *PI-P*spI (NEB), and the fused EGFP expression cassettes from the digested plasmids were integrated into the pRCS2-ocs-nptII vector for transformation (Chung et al. 2005).

For the CRISPR–Cas9-mediated knockout of *OsSWEET15*, two pairs of specific spacers (sequences listed in Supp. Table 1) of *OsSWEET15* were synthesized according to the rice CRISPR database (provided by Professor Lijia Qu) with additional *Bsa*I restriction site sequences at either end. The annealed double-stranded spacers were inserted into the *Bsa*I-digested pOs-sgRNA vector. A Gateway Cloning LR reaction (Invitrogen) of the resulting pOs-sgRNA constructs was performed with the destination vector pH-Ubi-Cas9-7 (Miao et al. 2013). All destination plasmids were confirmed by sequencing.

### Transient Expression of OsSWEET15–EGFP Fusion Protein in Rice Protoplast

Protoplasts were isolated from culms of etiolated rice seedlings and were subsequently transfected with plasmids pSAT6A-EGFP-OsSWEET15+CD3-1007 using a PEG mediated transfection method (Miao and Jiang, 2007; Ying et al., 2017). The plasmid CD3-1007 (Nelson et al., 2007) containing the plasma membrane protein AtPIP2;1 marked with mCherry (AtPIP2;1-mCherry fusion) were used to indicate the plasma membrane (Breia et al., 2020). pSAT6A-EGFP-N1+CD3-1007 were co-transfected in parallel as the negative control; and pSAT6A-EGFP-OsSWEET11 + CD3-1007 were co-transfected in parallel as the positive control. Detection of the fluorescence signal in the rice protoplasts was performed with a confocal laser scanning microscope (Leica SP5) after incubation of the protoplasts in the dark at room temperature for 12-15h. The excitation/emission wavelength of GFP is 488/506 to 538 nm, and the excitation/emission wavelength of mCherry fluorescent protein is 561/575 to 630 nm.

### Transforming and Screening Positive Transgenic Rice

Plasmids harboring pOsSWEET15::GUS, pOsSWEET15::GFP cassettes, and constructs for CRISPR-Cas9 gene editing were introduced into *Agrobacterium tumefaciens* strain EHA105 by electroporation, and positive agrobacteria were used to infect rice Nipponbare callus. The pOsSWEET15::GUS, pOsSWEET15::GFP, and CRISPR-Cas9 edited transgenic lines were initially screened based on hygromycin resistance. They were subsequently evaluated by PCR amplification with gene-specific primers P9/P10 (Supp. Table 1) and fragments were sequenced. Seedlings about 15d after germination of transgenic and WT rice as control were transplanted into a rice field in Nanjing (N:32°01’, E:118°51’, altitude: 18m.), China, in late April each year and grow in natural conditions with a light/dark regime around 14h/10h in early May.

### Quantitative Real-Time PCR Analyses of the Expression of *OsSWEET15* in Different Tissues of Rice

Total RNAs were extracted from rice tissues using the TRIzol reagent (Invitrogen), following the manufacturer’s protocol. Root, stem, and leaf samples were collected from 3-week-old hydroponic rice seedlings. Anthers were collected from rice plants approaching flowering. Caryopses were collected from spikelets 2, 3, 4, and 5 days after flowering (DAF). Quantitative real-time reverse transcription-PCR (qRT-PCR) amplification was performed with primers P11/P12 for the *OsSWEET15* fragment. P13/P14 for amplification of *OsActin* (LOC_Os03g50890) cDNA fragment (AB047313) and P15/16 for amplification of α-Tubulin (LOC_Os07g38730) cDNA fragment (AK104900) as housekeeping genes (Li et al., 2010). PCR products were confirmed by sequencing. Expression levels of *OsSWEET15* were normalized based on the expression levels of Actin or α-Tubulin with three biological replicates according to the 2^−ΔΔCT^ method (Livak and Schmittgen, 2001). All the primer sequences for amplification are listed in Supp. Table 1.

### Tissue Localization of OsSWEET15 Analyzed through Histochemical Staining and GFP Fluorescence Observation

Rice tissues from four different transgenic rice lines containing the pOsSWEET15 (2262 bp)::GUS expression cassette were collected and submerged into a GUS reaction mix, and were incubated at 37 °C overnight as described by Ma et al. (2017). Caryopses were collected at 5 DAF. Pollens were collected from flowering inflorescences. Root, leaf blade, and sheath samples were collected from 3-week-old rice seedlings. GUS-stained samples were observed under a stereomicroscope (Olympus MVX10) or a microscope (Olympus BX51). To obtain detailed expressional information of the protein, the GUS-stained tissues were rinsed and fixed in a FAA solution (formalin: glacial acetic acid: 50 % ethanol=1:1:18) for 24 h, and were then embedded in Epon812 (Sigma) as described by Coulter (1967). Fresh sections of pOsSWEET15::GFP expressing caryopses were observed using a confocal microscope (SP5; Leica).

### Transmission Electron Microscopy

The upper parts of caryopses of WT rice at 5 DAF were removed and the bottom parts were fixed in 2.5 % glutaraldehyde solution. They were further fixed in 1 % Osmium tetroxide after being rinsed with 0.1M PBS solution. After gradient ethanol dehydration, the samples were embedded in Epon812 and sliced longitudinally and transversely. The sliced samples were double-stained with uranyl acetate and lead citrate before being observed using a transmission electron microscope (Hitachi H-7650) at an accelerating voltage of 80 kV.

### Test of Pollen Germination

Pollens in stamens of *ossweet15* mutant lines and wild-type rice were scattered on Petri dishes containing a medium consisting of 5 % potato starch, 5g /L H_3_BO_3_, 15 % sucrose. They were incubated at 28 °C for one hour before being observed and photographed under a microscope (Olympus BX51).

### Observation of Embryo and Endosperm Development

Nucleate in the developing embryo sac of *ossweet15* mutant lines and WT control were sucked out by a pipette with 10 μl tips, they were placed on slides and stained with a DAPI solution in the dark at room temperature for 10 mins. DAPI fluorescence was observed with a Leica microscope (DM5000B).

## Abbreviations

CRISPR-Cas9: Clustered Regularly Interspaced Short Palindromic Repeats-Cas9
DAF: days after flowering
DAPI: 4′, 6-diamidino-2-phenylindole
ESR: embryo surrounding region
FRET: Förster resonance energy transfer
MCE: micropylar endosperm

## Supplemental Data

**Supplemental Figure S1**. OsSWEET15 expression in different tissues of rice (Nipponbare) indicated by GUS staining. OsSWEET15 was expressed in root, leaf blade, sheath, and stem. Samples were collected from 3-week-old rice seedlings. Bars in a =250 μm; in b =1 mm; in c=3 mm.

**Supplemental Figure S2**. Confirmation of CRISPR-Cas9 edited *OsSWEET15* mutant rice lines. DNA fragments of *OsSWEET15* in mutant lines were amplified with a high-fidelity polymerase and sequenced. Sequences and diagrams under the first-row sequence of each panel are CRISPR-Cas9 edited sequences.

**Supplemental Figure S3**. Comparison of *ossweet11* mutant rice lines and WT control (Nipponbare) grew in the year 2021 and statistical analyses of their 1000-grain weights.

**Supplemental Figure S4**. Comparison of predicted OsSWEET15 protein structures (right) of the mutants and WT control. Transmembrane helix prediction of *OsSWEET15* mutant and WT proteins was performed using TMHMM (http://www.cbs.dtu.dk/services/TMHMM/).

**Supplemental Figure S5**. Spike and caryopsis comparisons of *OsSWEET11/OsSWEET15* double gene mutant lines and WT rice control (Nipponbare). Upper panel: spikes of 3 double-gene mutant lines of (m11/15-15, m11/15-16, m11/15-21) (upper left) and WT (upper right). Lower panel: caryopses of the double gene mutants and WT. The spikelets of the double gene mutants either bore withered grains or were simply empty. Bar=0.3mm in the lower panel.

**Supplemental Table S1**. PCR primer pairs for cloning, checking and expression analyses or knockout of *OsSWEET15* in rice.

## ACKNOWLEDGMENT

We thank Professor Lijia Qu at Peking University for providing us with the CRISPR gene-editing tool.

## Parsed Citations

Arshad MS, Farooq M, Asch F, Krishna JSV, Prasad PVV, Siddique KHM (2017) Thermal stress impacts reproductive development and grain yield in rice. Plant Physiol Biochem 115:57–72

Bai AN, Lu XD, Li DQ, Liu JX, Liu CM (2015) NF–YB1-regulated expression of sucrose transporters in aleurone facilitates sugar loading to rice endosperm. Cell Res 25:1–5

Becraft PW (2001) Cell fate specification in the cereal endosperm. Semin Cell Dev Biol 12:387–394

Berger F (2003) Endosperm: The crossroad of seed development. Curr Opin Plant Biol 6: 42–50

Breia R, Conde A, Pimente D, Conde C, Fortes AM, Granell A, Gerós H (2020) VvSWEET7 Is a Mono- and Disaccharide Transporter Up-Regulated in Response to Botrytis cinerea Infection in Grape Berries. Front Plant Sci 10:1753

Briggs CL (1996) An ultrastructural study of the embryo/endosperm interface in the developing seeds of Solanum nigrum L. zygote to mid torpedo stage. Ann Bot 78:295–304

Chen LQ, Hou BH, Lalonde S, Takanaga H, Hartung ML, Qu XQ, Guo WJ, Kim JG, Underwood W, Chaudhuri B, et al (2010) Sugar transporters for intercellular exchange and nutrition of pathogens. Nature 468:527–532

Chen LQ, Qu XQ, Hou BH, Sosso D, Osorio S, Fernie AR, FrommerWB (2012) Sucrose efflux mediated by SWEET proteins as a key step for phloem transport. Science 335:207–211.

Chena LQ, Cheung LS, Feng L, Tanner W, Frommer WB (2015) Transport of sugars. Annu Rev Biochem 84:865–94

Chenb LQ, Lin IW, Qu XQ, Sosso D, McFarlane HE, Londoño A, Samuels AL, Frommer WB (2015) A cascade of sequentially expressed sucrose transporters in the seed coat and endosperm provides nutrition for the Arabidopsis embryo. Plant Cell 27: 607–619

Chu Z, Yuan M, Yao J, Ge X, Yuan B, Xu C, Li X, Fu B, Li Z, Bennetzen JL, et al (2006) Promoter mutations of an essential gene for pollen development result in disease resistance in rice. Genes Dev 20:1250–1255

Chung SM, Frankman EL and Tzfira T (2005)A versatile vector system for multiple gene expression in plants. Trends Plant Sci 10:357–361

Coulter HD (1967) Rapid and improved methods for embedding biological tissues in Epon 812 and Araldite 502. J Ultrastruct Res 20:346–355

Dumas C, Rogowsky P (2008) Fertilization and early seed formation. Fertilization and early seed formation. C R Biol 331:715–725

Dute RR, Peterson CM, Rushing AE (1989) Utrastructural changes of the egg apparatus associated with fertilization and proembryo development of soybean, Glycine max (Fabaceae). Ann Bot 64:123–136

Engell K (1989) Embryology of barley: Time course and analysis of controlled fertilization and early embryo formation based on serial sections. Nord J Bot 9:265–280

Eom JS, Chen LQ, Sosso D, Julius BT, Lin IW, Qu XQ, Braun DM, Frommer WB (2015) SWEETs, transporters for intracellular and intercellular sugar translocation. Curr Opin Plant Biol 25:53–62

Fei H, Yang Z, Lu Q, Wen X, Zhang Y, Zhang A, Lu C (2021) OsSWEET14 cooperates with OsSWEET11 to contribute to grain filling in rice. Plant Sci 306:110851.

Guo W-J, Nagy R, Chen H-Y, Pfrunder S, Yu Y-C, Santelia D, Frommer WB, Martinoia E (2014) SWEET17, a facilitative transporter, mediates fructose transport across the tonoplast of Arabidopsis roots and leaves. Plant Physiol164:777–89

Hehenberger E, Kradolfer D, Kohler C (2012) Endosperm cellularization defines an important developmental transition for embryo development. Dev 139:2031–2039

Higashiyama T, Takeuchi H (2015) The mechanism and key molecules involved in pollen tube guidance. Annu Rev Plant Biol 66:393–413

Itoh J, Nonomura K, Ikeda K, Yamaki S, Inukai Y, Yamagishi H, Kitano H, Nagato Y (2005) Rice plant development: from zygote to spikelet. Plant Cell Physiol 46:23–47

Jinek M, Chylinski K, Fonfara I, Hauer M, Doudna JA and Charpentier E. (2012) A programmable dual RNA-guided DNA endonuclease in adaptive bacterial immunity. Science 337:816–821

Kanno Y, Oikawa T, Chiba Y, Ishimaru Y, Shimizu T, Sano N, Koshiba T, Kamiya Y, Ueda M, Seo M (2016) AtSWEET13 and AtSWEET14 regulate gibberellin-mediated physiological processes. Nat Commun 7: 13245

Kawashima T and Goldberg RB (2009) The suspensor: not just suspending the embryo. Trends Plant Sci 15:23–30

Ko H-Y, Ho L-H, Neuhaus HE, Guo W-J (2021) Transporter SlSWEET15 unloads sucrose from phloem and seed coat for fruit and seed development in tomato. Plant Physiol kiab290.

Lafon-Placette C, Köhler C (2014) Embryo and endosperm, partners in seed development. Curr Opin Plant Biol 17:64–69

Livak KJ and Schmittgen TD (2001) Analysis of relative gene expression data using real-time quantitative PCR and the 2(-Delta Delta C(T)). Methods 25 402–408

Li J, Wu L, Foster R, Ruan YL (2017) Molecular regulation of sucrose catabolism and sugar transport for development, defence and phloem function. J Integr Plant Biol 59:322–335

Lin IW, Sosso D, Chen LQ, Gase K, Kim SG, Kessler D, Klinkenberg PM, Gorder MK, Hou BH, Qu XQ, et al (2014) Nectar secretion requires sucrose phosphate synthases and the sugar transporter SWEET9. Nature 508:546–549

Ma L, Zhang D, Miao Q, Yang J, Xuan Y, Hu Y (2017) Essential role of sugar transporter OsSWEET11 during the early stage of rice grain filling. Plant Cell Physiol 58:863–873

Mansfield SG, Briarty LG (1991) Early embryogenesis in Arabidopsis thaliana. II. The developing embryo. Can J Bot 69:461–476

Matsukura C, Saitoh T, Hirose T, Ohsugi R, Perata P, and Yamaguchi J (2000) Sugar uptake and transport in rice embryo. Expression of companion cell-specific sucrose transporter (OsSUT1) induced by sugar and light. Plant Physiol 124:85–93

Miao J, Guo D, Zhang J, Huang Q, Qin G, Zhang X, Wan J, Gu H, Qu LJ (2013) Targeted mutagenesis in rice using CRISPR-Cas system. Cell Res 23:1233–1236

MiaoYS, Jiang LW (2007) Transient expression of fluorescent fusion proteins in protoplasts of suspension cultured cells. Nat Protoc 2:2348–2353

Morii M, Sugihara A, Takehara S, Kanno Y, Kawai K, Hobo T, Hattori M, Yoshimura H, Seo M, Ueguchi-Tanaka M (2020) The dual function of OsSWEET3a as a gibberellin and glucose transporter is important for young shoot development in rice. Plant Cell Physiol 61:1935–1945

Morley-Smith ER, Pike MJ, Findlay K, Kockenberger W, Hill LM, Smith AM, Rawsthorne S (2008) The transport of sugars to developing embryos is not via the bulk endosperm in oilseed rape seeds. Plant Physiol 147:2121–2130

Nelson BK, Cai X and Nebenfuhr A (2007) A multicolored set of in vivo organelle markers for co-localization studies in Arabidopsis and other plants. Plant J 51: 1126–1136

Oliva R, Ji C, Atienza-Grande G, Huguet-Tapia JC, Perez-Quintero A, Li T, Eom JS, Li C, Nguyen H, Liu B, et al (2019) Broadspectrum resistance to bacterial blight in rice using genome editing. Nat Biotechnol 37:1344–1350

Olsen OA (2001) ENDOSPERM DEVELOPMENT: Cellularization and cell fate specification. Annu Rev Plant Physiol Plant Mol Biol 52:233–267

Olsen OA (2004) Nuclear Endosperm Development in Cereals and Arabidopsis thaliana. Plant Cell 16:S214–S227

Opsahl-Ferstad HG, Le Deunff E, Dumas C, Rogowsky PM (1997) ZmEsr, a novel endosperm-specific gene expressed in a restricted region around the maize embryo. Plant J 12:235–246

Schel JHN, Kieft H and van Lammeren AAM (1984) Interaction between embryo and endosperm during early developmental stages of maize caryopses (Zea mays). Can J Bot 62:2842–2853

Sosso D, Luo D, Li QB, Sasse J, Yang J, Gendrot G, Suzuki M, Koch KE, McCarty DR, Chourey PS, et al (2015) Seed filling in domesticated maize and rice depends on SWEET-mediated hexose transport. Nat Genet 47:1489–1493

Suzuki K, Miyake H, Taniguchi T, Maeda E (1993) Ultrastructural Studies on Rice Globular Embryos with Emphasis on Epidermis Initiation. JPN J Crop Sci 62:116–125

Suzuki K, Miyake H, Taniguchi T, Maeda E (2000) Cellularization of the Free Nuclear Endosperm in Rice Caryopsis Revealed by Light and Electron Microscopy. Plant Prod Sci 3:446–458

Wang S, Yokosho K, Guo R, Whelan J, Ruan Y-L, Ma JF, Shou H (2019) The soybean sugar transporter GmSWEET15 mediates sucrose export from endosperm to early embryo. Plant Physiol 180:2133–2141

Wang X, Liu X, Hu Z, Bao S, Xia H, Feng B, Ma L, Zhao G, Zhang D, Hu Y. Essentiality for rice fertility and alternative splicing of OsSUT1. Plant Sci. 2021. doi.org/10.1016/j.plantsci.2021.111065

Wua X, Liu J, Li D, Liu CM (2016) Rice caryopsis development I: Dynamic changes in different cell layers. J Integr Plant Biol 58:772–785

Wub X, Liu J, Li D, Liu CM (2016) Rice caryopsis development II: Dynamic changes in the endosperm. J Integr Plant Biol 58:786–798

Xu Y, Yang J, Wang Y, Wang J, Yu Y, Long Y, Wang Y, Zhang H, Ren Y, Chen J, et al (2017) OsCNGC13 promotes seed-setting rate by facilitating pollen tube growth in stylar tissues. PLoS Genet 13:e1006906.

Ying Y, Yue W, Wang S, Li S, Wang M, Zhao Y, Wang C, Mao C, Whelan J, Shou H (2017) Two h-Type Thioredoxins Interact with the E2 Ubiquitin Conjugase PHO2 to Fine-Tune Phosphate Homeostasis in Rice. Plant Physiol 173:812–824

Yang B, Sugio A, White FF (2006) Os8N3 is a host disease-susceptibility gene for bacterial blight of rice. Proc Natl Acad Sci USA 103:10503–10508

Yang J, Luo D, Yang B, Frommer WB, Eom JS (2018) SWEET11 and 15 as key players in seed filling in rice. New Phytol 218:604–615

Zhang W-H, Zhou Y, Dibley K, Tyerman S, Furbank R, Patrick J (2007) Nutrient loading of developing seeds. Funct Plant Biol 34:314–331

Zhou SR, Yin LL, Xue HW (2013) Functional genomics based understanding of rice endosperm development. Curr Opin Plant Biol 16:236–246

